# Combinatorial interactions between viral proteins expand the functional landscape of the viral proteome

**DOI:** 10.1101/2021.04.07.438767

**Authors:** Liping Wang, Huang Tan, Laura Medina-Puche, Mengshi Wu, Borja Garnelo Gómez, Man Gao, Chaonan Shi, Tamara Jimenez-Góngora, Pengfei Fan, Xue Ding, Dan Zhang, Ding Yi, Tábata Rosas-Diaz, Yujing Liu, Emmanuel Aguilar, Xing Fu, Rosa Lozano-Durán

## Abstract

As intracellular parasites, viruses need to manipulate the molecular machinery of their host cells in order to enable their own replication and spread. This manipulation is based on the activity of virus-encoded proteins. The reduced size of viral genomes imposes restrictions in coding capacity; how the action of the limited number of viral proteins results in the massive cell reprogramming observed during the viral infection is a long-standing conundrum in virology. In this work, we explore the hypothesis that combinatorial interactions expand the multifunctionality of viral proteins, which may exert different activities individually and when in combination, physical or functional. We show that the proteins encoded by a plant-infecting DNA virus physically associate with one another in an intricate network. Our results further demonstrate that these interactions can modify the subcellular localization of the viral proteins involved, and that co-expressed interacting viral proteins can exert novel biological functions *in planta* that go beyond the sum of their individual functions. Based on this, we propose a model in which combinatorial physical and functional interactions between viral proteins enlarge the functional landscape of the viral proteome, which underscores the importance of studying the role of viral proteins in the context of the infection.

## INTRODUCTION

Viruses are intracellular parasites that need to subvert the host cell in order to enable their replication and ensure viral spread. For this purpose, viruses co-opt the cell molecular machinery, modulating or redirecting its functions; as a result, infected cells undergo dramatic molecular changes, including heavy transcriptional reprogramming, concomitant to the proliferation of the virus.

Most viruses have small genomes, which imposes limitations in coding capacity, with viral proteins frequently exhibiting small size, and with their numbers per viral genome ranging from a few (<10) to a few dozen (Supplementary figure 1). Viral proteins are known to be multifunctional, and have been suggested to target hubs in the proteomes of their host cells (Brito and Pinney, 2017; Dyer et al., 2011; King et al., 2018; Zheng et al., 2014), hence maximizing the impact of the viral-host protein-protein interactions; nevertheless, how a limited repertoire of small viral proteins can lead to the drastic cellular changes observed during the viral infection remains puzzling. Upon viral invasion, virus-encoded proteins are produced in large amounts in the infected cells, where they co-exist. Therefore, physical or functional interactions among viral proteins might have evolved as a potential mechanism to expand the virus-host functional interface, increasing the number of potential targets in the host cell and/or synergistically modulating the cellular environment. Interestingly, examples of interactions between viral proteins have been recently documented for both animal and plant viruses (e.g. (Ashford et al., 2016; Bragg and Jackson, 2004; Calderwood et al., 2007; Dao et al., 2020; DeBlasio et al., 2018; Fossum et al., 2009; Hagen et al., 2014; Leastro et al., 2018; Lee et al., 2011; Li et al., 2020; Li et al., 2021; Liu et al., 2010; Loureiro et al., 2012; Nobre et al., 2019; Rozen et al., 2008; Stellberger et al., 2010; Uetz et al., 2006; Varasteh Moradi et al., 2020; von Brunn et al., 2007); see VirHostNet 2.0, http://virhostnet.prabi.fr/, Guirimand et al., 2015); some of these interactions have been proposed to contribute to viral genome replication and virion assembly. However, the hypothesis that the combination of individual virus-encoded proteins might result in the acquisition of novel functions still lacks experimental support.

Here, we use the plant DNA virus *Tomato yellow leaf curl virus* (TYLCV; Fam. *Geminiviridae*) to test the idea that combinatorial interactions among viral proteins exist and may underlie an expansion of the functional landscape of the viral proteome. TYLCV encodes six proteins (C1/Rep, C2, C3, C4, V2, and CP); local infection by TYLCV in the experimental host *Nicotiana benthamiana* results in heavy transcriptional reprogramming, with 11,850 differentially expressed genes (DEGs) detected at 6 days post-inoculation (dpi) (Wu et al., 2019). Although a limited number of viral protein-protein interactions have been described for this virus (Hallan and Gafni, 2001; Settlage et al., 2005; Wang et al., 2020; Wang et al., 2017b; Zhao et al., 2018; Zhao et al., 2020), the intra-viral interactome has not been systematically explored, and the functional impact of these interactions remains elusive. Our results show that viral proteins form complexes in the context of the viral infection, displaying a high degree of intra-viral connectivity. As proof-of-concept, we focus on the pair formed by C2 and CP, since the presence of the latter is required and sufficient to shift the subcellular localization of the former; our data indicate that the combination of C2 and CP results in drastic transcriptional reprogramming in the host plant, which goes beyond the sum of the effects of each of the individual proteins.

## RESULTS AND DISCUSSION

### Viral proteins form complexes in the host cell

In order to test whether virus-encoded proteins associate with one another, we employed a number of protein-protein interaction methods, namely yeast two-hybrid (Y2H), *in planta* co-immunoprecipitation (co-IP), bimolecular fluorescence complementation (BiFC), and split-luciferase assays. Several viral protein-protein interactions were identified in yeast (Figure 1A; Supplementary figure 2); the number of associations between viral proteins found in co-IP was higher, and some of them were dependent on the presence of the virus (Figure 1B; Supplementary figure 3; Supplementary figure 4). These interactions were further confirmed in BiFC and split-luciferase experiments (Figure 1C, D). BiFC indicates that most of the detected interactions occur in the nucleus (Figure 1C; Supplemental figure 5; additional patterns of interactions observed by BiFC can be found in Supplementary figure 6). A summary of all detected interactions between viral proteins is shown in Figure 1E; all viral proteins were found to interact with one another, including self-interactions, by at least two independent methods. Importantly, some of these interactions could also be detected in unbiased affinity purification followed by mass spectrometry (AP-MS) experiments with GFP-tagged versions of the viral proteins expressed in infected *N. benthamiana* cells (Wang et al., 2017a), indicating that viral proteins physically associate with one another in the context of the infection (Supplementary table 1).

**Figure 1.**
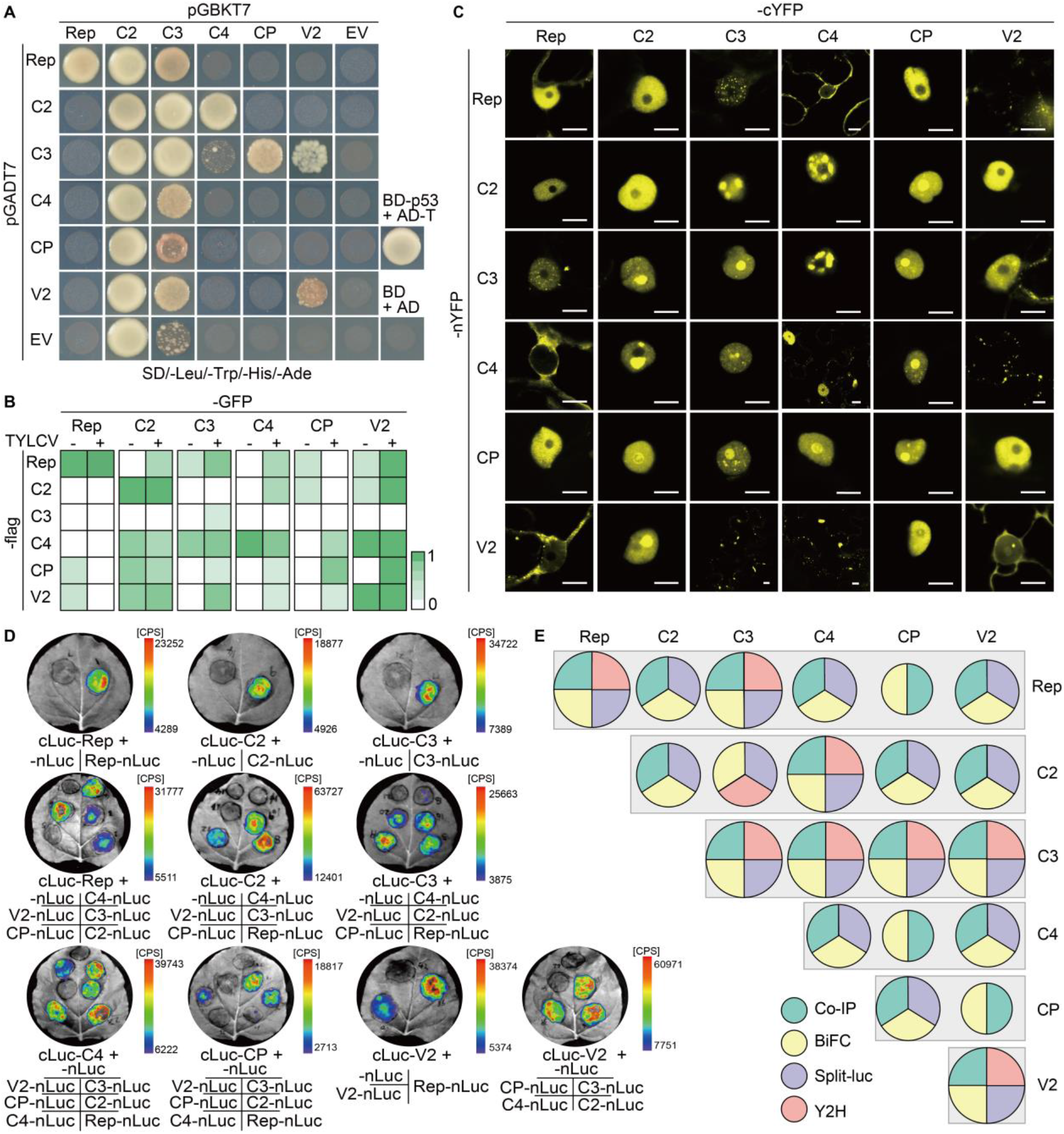
The proteins encoded by the plant DNA virus *Tomato yellow leaf curl virus* associate with one another in the plant cell. (**A**) Viral protein-protein interactions detected in yeast two-hybrid. The minimal synthetic defined (SD) medium without leucine (Leu), tryptophan (Trp), histidine (His), and adenine (Ade) was used to select positive interactions; SD without Leu and Trp was used to select co-transformants (Supplementary figure 2). The interaction between the SV40 large T antigen (T) and the tumor suppressor p53 is a positive control. AD: activation domain; BD: binding domain. This experiment was repeated three times with similar results. (**B**) Summary of viral protein-protein interactions detected by co-immunoprecipitation (co-IP) in the absence or presence of *Tomato yellow leaf curl virus* (TYLCV). These experiments were repeated at least three times; the colour scale represents the percentage of positive interaction results among all replicates, with 1=100%. The original co-IP blots are shown in Supplementary figure 2 (in the absence of TYLCV) and Supplementary figure 3 (in the presence of TYLCV). (**C**) Viral protein-protein interactions detected by bimolecular fluorescence complementation (BiFC) in *N. benthamiana* leaves. nYFP: N-terminal half of the YFP; cYFP: C-terminal half of the YFP. Images were taken at 2 days post-infiltration (dpi). Scale bar = 10 µm. This experiment was repeated at least four times with similar results; combination with Hoechst staining and negative controls can be found in Supplementary figure 5. Additional images are shown in Supplementary figure 6. (**D**) Viral protein-protein interactions detected by split-luciferase assay in *N. benthamiana* leaves. nLuc: N-terminal part of the luciferase protein; cLuc: C-terminal part of the luciferase protein. Images were taken at 2 dpi. The colour scale represents the intensity of the interaction in counts per second (CPS). This experiment was repeated three times with similar results. (**E**) Summary of the intra-viral protein-protein interactions identified in A-D. Different colours represent different methods, as indicated; circle size indicates the number of the methods in which a positive interaction was detected.

### The viral CP is required and sufficient to modify the subcellular localization of the virus-encoded C2 protein

Although the proteins encoded by TYLCV display specific localizations in the plant cell, all of them, with the exception of C4, can be consistently found in the nucleus in basal conditions (Figure 2A). Interestingly, in the presence of the virus, several viral proteins fused to GFP, namely C2, C3, C4, and CP, experienced obvious changes in their subcellular distribution (Figure 2A). These changes had been previously observed for C4 and CP; while in the case of C4, Rep alone can trigger its re-localization from the plasma membrane to chloroplasts (Medina-Puche et al., 2020), no individual protein was sufficient to modify the subnuclear pattern of CP (Wang et al., 2017b). In the absence of the virus, C2-GFP appears evenly distributed in the nucleoplasm and is excluded from the nucleolus, where it strongly accumulates when the virus is present; based on this gain of localization, we reasoned that C2 might perform additional functions in the context of the infection. Binary combinations with the virus-encoded proteins fused to RFP indicated that CP is sufficient to induce the localization of C2-GFP in the nucleolus (Figure 2B), where both proteins interact (Figure 1C); this was further confirmed by co-expression with the untagged version of the protein (Figure 2C; Supplementary figure 7A). Of note, only C2-GFP, but not GFP-C2, can be re-localized by CP (Figure 2C). Importantly, removal of the start codon and alternative transcription initiation sites in the CP ORF rendered the virus unable to re-localize C2 to the nucleolus (Figure 2D; Supplementary figure 7B), indicating that CP is not only sufficient, but also required for this change to occur in infected cells.

**Figure 2.**
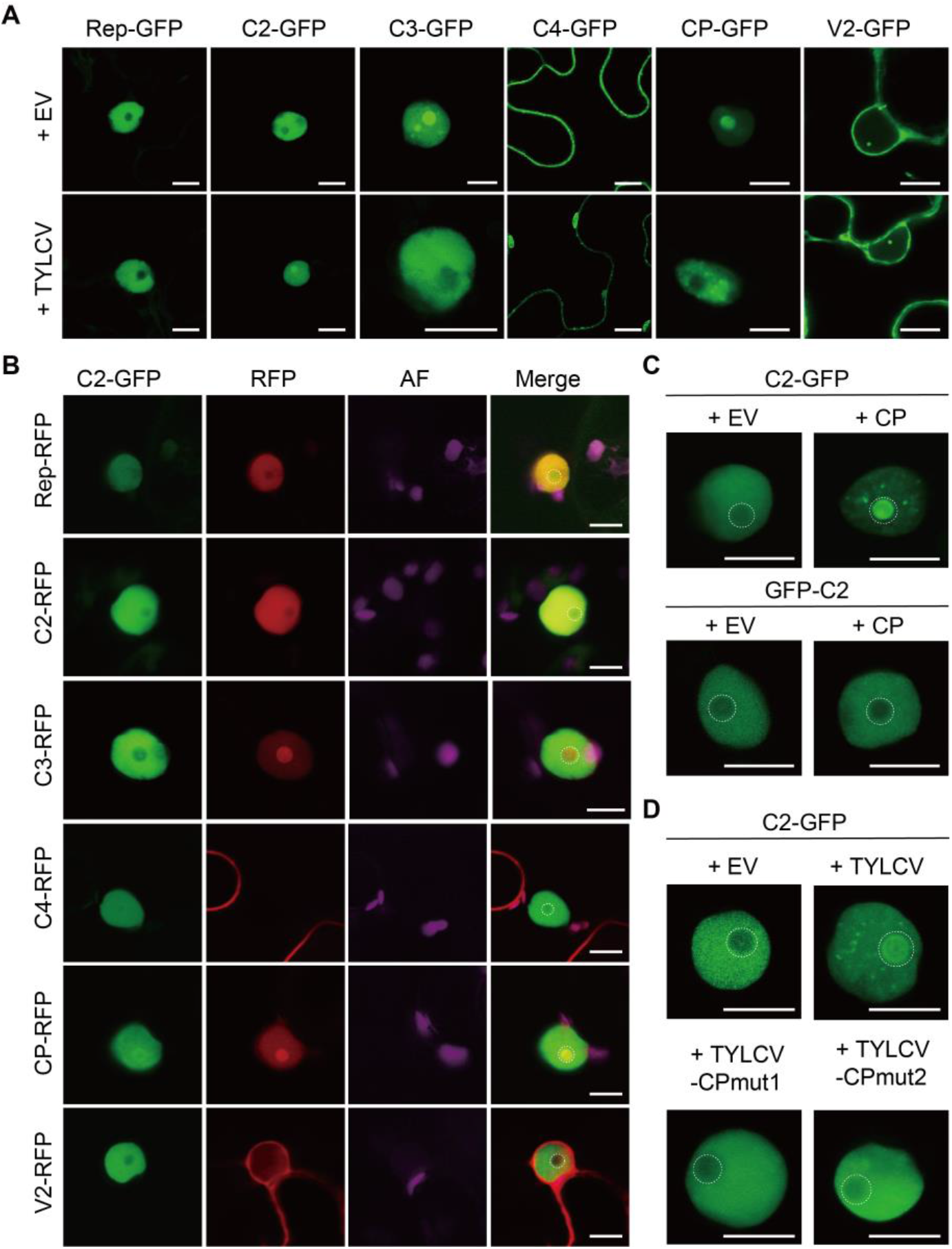
CP is required and sufficient to change the subnuclear localization of C2. (**A**) Subcellular localization of the TYLCV-encoded proteins fused to GFP expressed alone (+EV; co-transformed with an empty vector control) or in the context of the viral infection (+TYLCV; co-transformed with a TYLCV infectious clone) in *N. benthamiana* leaves at 2 days post infiltration (dpi). Scale bar = 10 µm. EV: empty vector. (**B**) Subcellular localization of C2-GFP expressed alone or co-expressed with each of the viral proteins fused to RFP in *N. benthamiana* leaves at 2 dpi. Scale bar = 10 µm. AF: Autofluorescence. (**C**). Subcellular localization of C2-GFP or GFP-C2 when expressed alone (+EV) or co-expressed with CP (+CP) in *N. benthamiana* leaves at 2 dpi. The accumulation of the CP transcript is shown in Supplementary figure 7A. Scale bar = 10 µm. EV: empty vector. (**D**). Subcellular localization of C2-GFP when expressed alone (+EV) or in the context of the infection by the WT TYLCV virus (+TYLCV) or mutated versions unable to produce CP (+TYLCV-CPmut1; +TYLCV-CPmut2) in *N. benthamiana* leaves at 2 dpi. Viral accumulation is shown in Supplementary figure 7B. For details on TYLCV-CPmut1 and TYLCV-CPmut2, see Materials and Methods. Scale bar = 10 µm. EV: empty vector.

### The C2/CP module specifically modifies the host transcriptome and modulates plant defence

With the purpose of assessing if the functional landscape of C2 might be expanded when in the presence of CP, and considering that the C2 protein from geminiviruses has been previously described to impact host gene expression (Caracuel et al., 2012; Liu et al., 2014; Rosas-Diaz et al., 2016; Soitamo et al., 2012; Trinks et al., 2005; Yang et al., 2013), we decided to investigate the transcriptional changes triggered by C2 in the presence or absence of CP. To this aim, we expressed C2, CP, or C2+CP in *N. benthamiana* leaves and determined the resulting changes in the plant transcriptome by RNA-seq. As shown in Figure 3A, C2 alone caused the differential expression of 211 genes, while expression of CP did not significantly affect the plant transcriptional landscape; simultaneous expression of C2 and CP resulted in a moderate increase in the number of differentially expressed genes (DEGs) to 263 (Figure 3A; Supplementary figure 8A, B; validation of the RNA-seq results is presented in Supplementary figure 8C; Supplementary table 2). Strikingly, however, the identity and behavior of DEGs was dramatically changed by the presence of CP (Figure 3B, C), indicating that C2 and CP have a synergistic effect on the host transcriptome. Functional enrichment analysis unveiled that addition of CP indeed shifted the functional gene ontology (GO) categories transcriptionally reprogrammed by C2, and that certain categories appear as statistically over-represented in the subset of down-regulated genes only when both viral proteins are simultaneously expressed (Figure 3D, E; Supplementary table 3). To investigate the relevance of the re-localization of C2 (Figure 2C) for this effect, we selected DEGs specifically affected by the co-expression of C2 and CP, and tested the ability of C2-GFP (which re-localizes in the presence of CP) or GFP-C2 (which does not re-localize in the presence of CP) to affect their expression when combined with CP. As shown in Figure 3F and Supplementary Figure 8D, only C2+CP and C2-GFP+CP, but not GFP-C2+CP, affect the expression of the selected genes compared with C2, C2-GFP, or GFP-C2, respectively. This result suggests that the modification in subcellular localization of C2 mediated by CP is required for the impact of the combination of these proteins on gene expression. Stress-related GO functional categories are over-represented in the subsets of C2-triggered DEGs, but disappear when CP is present (Figure 3D), suggesting that the effect of C2 on the plant response to stress might change upon co-expression of CP. With the aim to test this idea, we subjected *N. benthamiana* tissue expressing C2, CP, C2+CP, or ß-glucuronidase (GUS) as a negative control to inoculation with the plant pathogenic bacterium *Pseudomonas syringae* pv. *tomato* DC3000 *ΔhopQ1-1*. Expression of C2 rendered the plant more resistant to the bacteria, while expression of CP did not impact bacterial multiplication; simultaneous expression of C2 and CP, however, led to a mild decrease in bacterial load, statistically different from the one caused by C2 alone (Figure 3G). These results demonstrate that the presence of CP modulates the impact of C2 on the response to this biotic stress.

**Figure 3.**
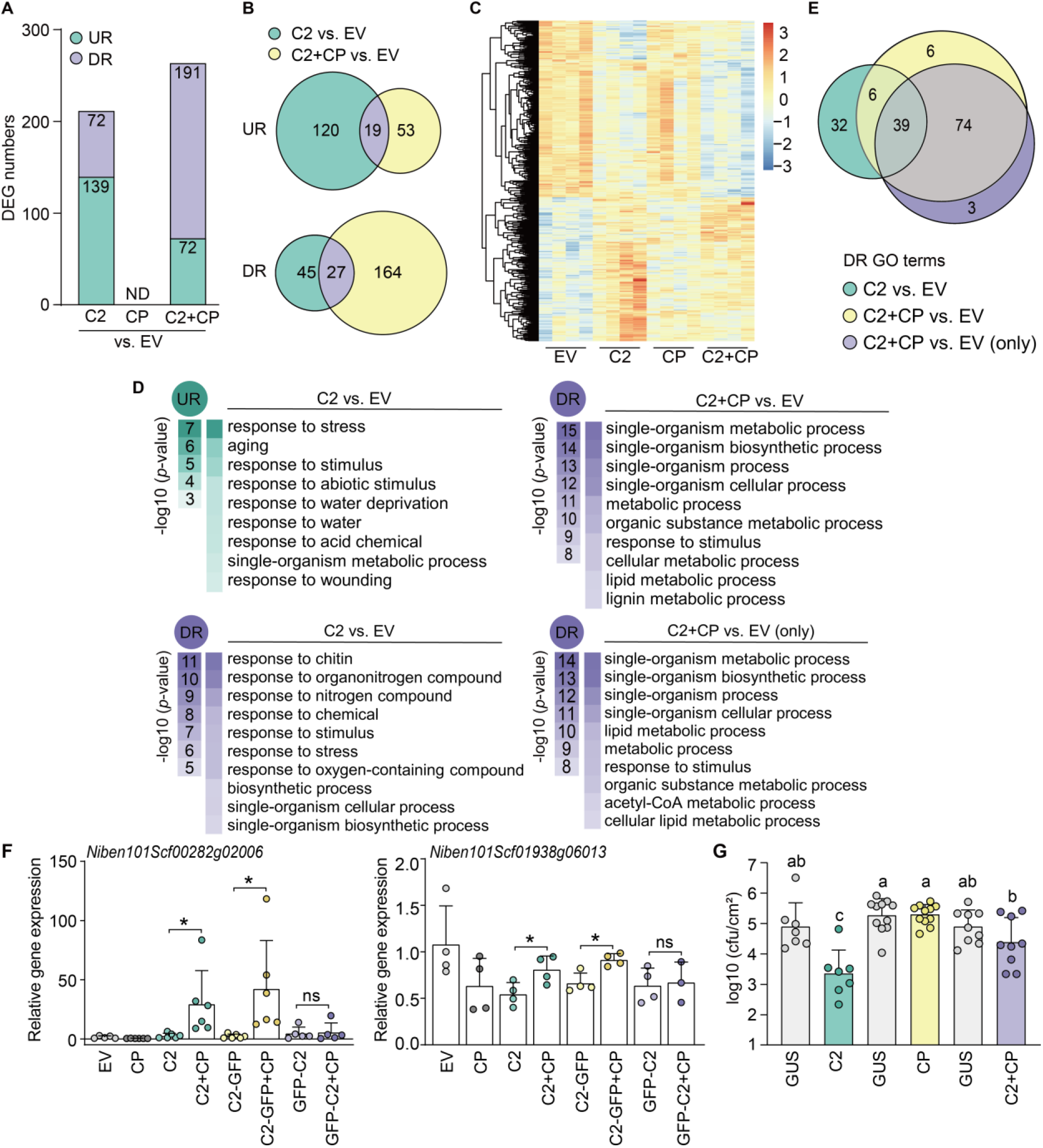
C2 and CP functionally interact *in planta* and modify the transcriptome of *N. benthamiana* in an interdependent manner. (**A**) Number of differentially expressed genes (DEGs) upon expression of C2, CP, or C2+CP in *N. benthamiana* leaves. UR: up-regulated; DR: down-regulated; ND: not detected; EV: empty vector. Full lists can be found in Supplementary table 2. (**B**) Venn diagram of DEGs upon expression of C2 or C2+CP in *N. benthamiana*. UR: up-regulated; DR: down-regulated; EV: empty vector. (**C**) Heatmap with hierarchical clustering from samples in (A). The colour scale indicates the Z-score. EV: empty vector. (**D**) Functional enrichment analysis of up-regulated (UR) or down-regulated (DR) genes in the indicated samples. Gene Ontology (GO) categories from the Biological Process ontology enriched with a *p-*value<0.01 (up to top 10) are shown; functional enrichment was performed using the orthologues in *Arabidopsis thaliana*. “C2+CP vs. EV (only)” denotes the subset of genes that are down-regulated in this sample only, and not in the samples expressing the viral proteins separately. The colour scale indicates the -log10 (*p*-value), showing the significance of GO term enrichment. EV: empty vector. For a full list, see Supplementary table 3. (**E**) Venn diagram of the GO terms (Biological Process ontology) over-represented in the subsets of down-regulated genes (*p*-value<0.01) in the different samples. DR: down-regulated; EV: empty vector. For a full list, see Supplementary table 3. (**F**) Expression of selected DEGs upon transient expression of C2, C2-GFP, or GFP-C2 in the presence and absence of CP in *N. benthamiana* leaves measured by qRT-PCR. The samples expressing CP or empty vector (EV) are used as control. Expression values are the mean of at least three biological replicates. Error bars represent SD. Asterisks indicate a statistically significant difference (*: *p*<0.05, **: *p*<0.01) according to a two-tailed comparison t-test. *NbACT* was used as the normalizer. (**G**) *Pseudomonas syringae* pv *tomato* DC3000 *ΔhopQ1-1* growth in *N. benthamiana* leaves expressing C2, CP, C2+CP, or ß-glucuronidase (GUS) as negative control. Values are the mean of more than six biological replicates. Error bars represent SD. Letters indicate a statistically significant difference (*p*<0.05) according to one-way ANOVA followed by post-hoc Tukey test. Experiments were repeated three times with similar results.

Next, we investigated the contribution of C2 and CP to the virus-induced transcriptional reprogramming in the context of the viral infection. We reasoned that, if C2 and CP together affect the transcriptional landscape of the host in a different manner than C2 or CP alone, then the transcriptional changes triggered by mutated versions of the virus unable to produce either C2 or CP should present overlapping differences compared to the changes triggered by the wild-type (WT) virus. Following this rationale, we compared the transcriptome of *N. benthamiana* leaves infected with the WT virus or mutated versions unable to produce C2 (TYLCV-C2mut) or CP (TYLCV-CPmut1), with respect to the empty vector (EV) control (Figure 4A) or to the WT virus (Figure 4B). As expected, both point mutants were unable to establish a full systemic infection, indicating that the corresponding viral proteins are most likely not produced from the mutated genes (Supplemental figure 9A, B). Of note, although the CP null mutant (TYLCV-CPmut1) replicated to lower levels, no significant changes in the accumulation of viral transcripts were detected among these viral variants in local infection assays (Supplementary figure 7B; Supplementary figure 9C-F). Importantly, and despite the fact that expression of CP alone did not result in detectable transcriptional changes, mutation of CP in the viral genome led to the differential expression of 3,256 genes when compared to the WT infection, supporting the notion that CP modulates host gene expression in combination, physical or functional, with other viral proteins; remarkably, 2,591 of these DEGs (79.5%) overlapped with those caused by the loss of C2 (Figure 4C; Figure 4D; Supplementary figure 9G; Supplementary table 2; validation of the RNA-seq results is presented in Supplementary figure 9H), indicating that C2 and CP cooperatively mediate changes in host gene expression during the infection. Functional categories over-represented among the up-regulated genes in the presence of the WT virus appear as down-regulated in the subset of DEGs commonly triggered by the C2-and CP-deficient viruses compared to the WT version (Figure 4E; Figure S10; Supplementary tables 4 and 5), suggesting that the C2/CP module is responsible for the transcriptional changes of genes associated to these GO terms. A complete overview of the functional enrichment in the different subsets of DEGs can be found in Supplementary figure 10 and Supplementary table 5.

**Figure 4.**
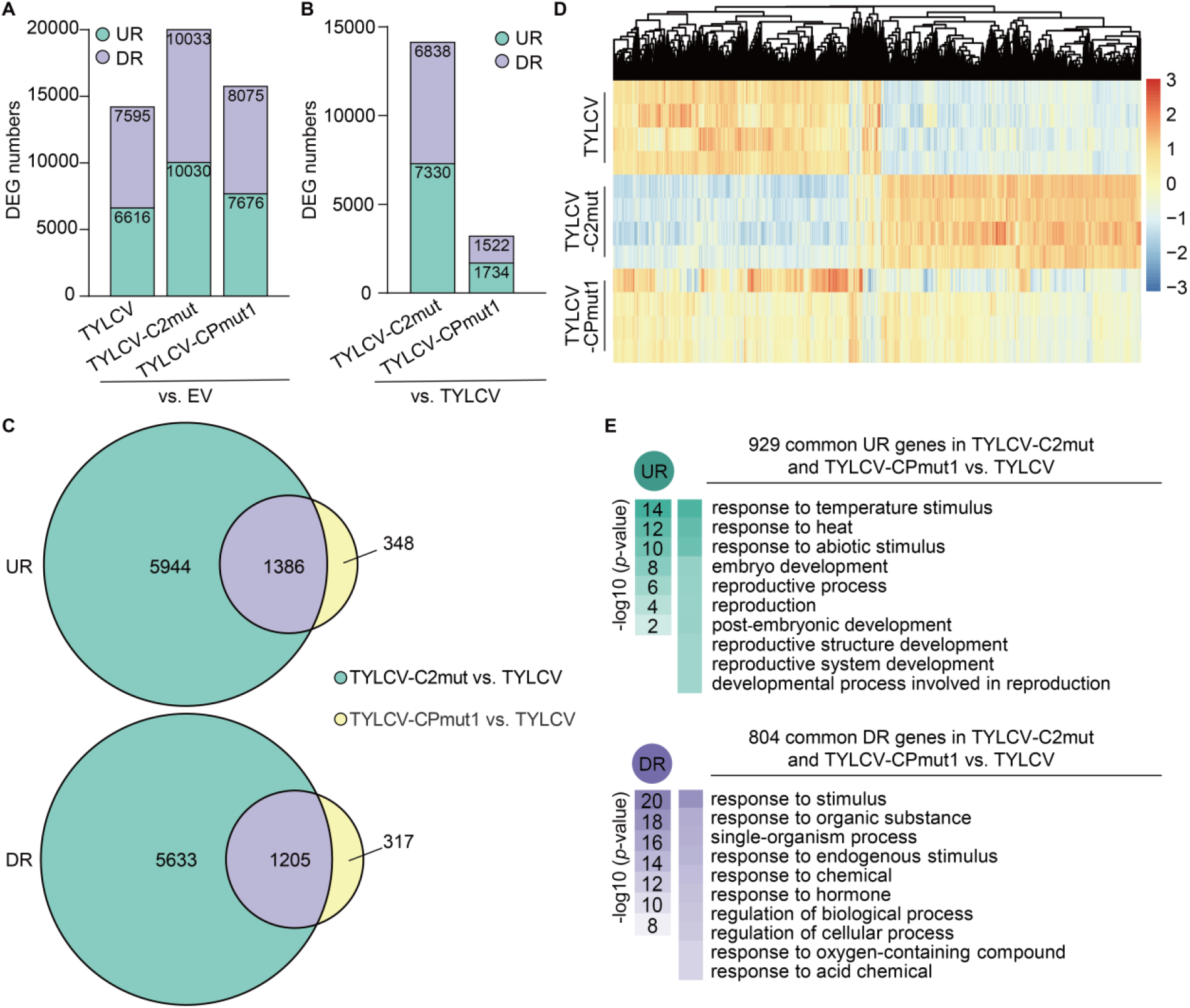
C2 and CP functionally interact *in planta* in the context of the viral infection. (**A** and **B**) Number of differentially expressed genes (DEGs) upon infection by TYLCV WT or C2-null or CP-null mutant variants (TYLCV-C2mut and TYLCV-CPmut1, respectively) in *N. benthamiana* leaves compared to the empty vector control (A), or to TYLCV WT (B). UR: up-regulated; DR: down-regulated; EV: empty vector. Full lists can be found in Supplementary table 2. (**C**) Venn diagrams of DEGs upon infection by TYLCV C2-null and TYLCV CP-null mutants (TYLCV-C2mut and TYLCV-CPmut1, respectively) compared to TYLCV WT. UR: up-regulated; DR: down-regulated. (**D**) Heatmap with hierarchical clustering from (A). The colour scale indicates the Z-score. (**E**) Functional enrichment analysis of the subsets of up-regulated (UR) or down-regulated (DR) genes in the indicated samples. Gene Ontology (GO) categories from the Biological Process ontology enriched with a *p-*value<0.01 (up to top 10) are shown; functional enrichment was performed using the orthologues in *Arabidopsis thaliana*. The colour scale indicates the -log10 (*p*-value), showing the significance of GO term enrichment. For a full list, see Supplementary table 4.

Taken together, our results demonstrate that TYLCV proteins form an intricate network of interactions that potentially vastly increase the complexity of the virus-host interface, and that viral proteins can exert additional functions when in combination. Given that intra-viral protein-protein interactions have been reported for viruses belonging to independently evolved families and infecting hosts belonging to different kingdoms of life, we propose that this might be an evolutionary strategy of viruses, which would call for a reconsideration of our approaches to the study of virus-host interactions.

## MATERIALS AND METHODS

### Plant material

*Nicotiana benthamiana* plants were grown in a controlled growth chamber in long-day conditions (16 h light/8 h dark) at 25°C.

### Bacterial strains and growth conditions

*Agrobacterium tumefaciens* strain GV3101 harbouring the corresponding binary vectors were liquid-cultured in LB medium (1% tryptone, 0.5% yeast extract, and 1% NaCl) with the appropriate antibiotics at 28°C overnight. *P. syringae* pv. *tomato* DC3000 *ΔhopQ1-1* (Rufian et al., 2018) was cultured on solid LB medium (1% tryptone, 0.5% yeast extract, and 0.5% NaCl) with the appropriate antibiotics at 28°C overnight.

### Plasmids and cloning

Open reading frames (ORFs, corresponding to Rep, C2, C3, C4, V2, and CP) from TYLCV (GenBank accession number AJ489258) were cloned in pENTR/D-TOPO (Thermo Scientific) without a stop codon (Wang et al., 2017a). The binary constructs to express viral proteins without tag, tagged with Cter-GFP, Nter-GFP, Cter-FLAG, or Cter-RFP, were generated by Gateway-cloning (LR reaction, Thermo Scientific) the TYLCV ORFs from pENTR/D-TOPO into pGWB2 (Nakagawa et al., 2007a), pGWB5 (Nakagawa et al., 2007a), pGWB6 (Nakagawa et al., 2007a), pGWB511 (Nakagawa et al., 2007b), and pGWB554 (Nakagawa et al., 2007b), respectively, with the exception of the construct to express C4-RFP, which was generated by Gateway-cloning the C4 ORF into pB7RWG2.0 (Karimi et al., 2002). For biomolecular fluorescence complementation assays (BiFC), the TYLCV ORFs were Gateway-cloned into pGTQL1211YN and pGTQL1221YC (Lu et al., 2010). For yeast two-hybrid assays (Y2H), pGBKT7 and pGADT7 (Clontech) were digested with *EcoR*I and *Pst*I or *EcoR*I and *BamH*I, respectively, and the PCR-amplified Rep, C2, C3, C4, V2, and CP ORFs were in-fused to the C-terminus of the GAL4 DNA-binding domain (in pGBKT7) and the C-terminus of GAL4 activation domain (in pGADT7) with ClonExpress® II One Step Cloning Kit (Vazyme). The binary constructs for split-luciferase complementation imaging assay were generated by Gateway cloning the TYLCV ORFs into pGWB-nLuc and pGWB-cLuc (Wang et al., 2019). The TYLCV infectious clone has been previously described (Rosas-Diaz et al., 2018). Using the wild-type (WT) infectious clone as template, the TYLCV C2 null mutant (TYLCV-C2mut), carrying a C-to-G mutation in the 14th nucleotide of the C2 ORF, was generated, converting the fifth codon (encoding a serine) to a stop codon, with the Quick Change Lightning Site-Directed Mutagenesis Kit (Agilent Technologies, Cat #210518). Similarly, the TYLCV CP null mutant 1 (TYLCV-CPmut1), carrying a C-to-A mutation in the fourth nucleotide of the CP ORF, was generated, converting the second codon (encoding a serine) to a stop codon. The TYLCV-CPmut2 infectious clone, containing two premature stop codons in positions 2 and 15 and in which the nine potential alternative starting sites (ATG) have been removed, was synthesized. In both cases, the mutations in the CP ORF do not affect the overlapping V2 ORF. All primers and plasmids used for cloning are summarized in Supplementary tables 6 and 7, respectively.

### Agrobacterium-mediated transient gene expression in *N. benthamiana*

Transient expression assays were performed as previously described (Wang et al., 2017a) with minor modifications. In brief, all binary plasmids were transformed into *A. tumefaciens* strain GV3101; *A. tumefaciens* clones carrying the constructs of interest were liquid-cultured in LB with the appropriate antibiotics at 28°C overnight. Bacterial cultures were collected by centrifugation at 4,000 x g for 10 min and resuspended in the infiltration buffer (10 mM MgCl_2_, 10 mM MES pH 5.6, 150 μM acetosyringone) to an OD_600_ = 0.2-0.5. Next, bacterial suspensions were incubated at room temperature in the dark for 2-4 hours before infiltration into the abaxial side of 4-week-old *N. benthamiana* leaves with a 1 mL needleless syringe. For experiments that required co-infiltration, the *Agrobacterium* suspensions carrying different constructs were mixed at 1:1 ratio before infiltration.

### Protein extraction and immunoprecipitation assays

Fully expanded young leaves of 4-week-old *N. benthamiana* plants were co-infiltrated with *A. tumefaciens* carrying constructs to express Rep-, C2-, C3-, C4-, CP-, and V2-flag, with Rep-, C2-, C3-, C4-, CP-or V2-GFP. To analyze these protein-protein interactions in the context of the viral infection, *A. tumefaciens* carrying the infectious TYLCV clone were co-infiltrated in the respective experiments. Two days after infiltration, 0.7-1 g of infiltrated *N. benthamiana* leaves were harvested. Protein extraction, co-immunoprecipitation (co-IP), and western blot were performed as previously described (Macho et al., 2014). For western blot, the following primary and secondary antibodies were used: mouse anti-green fluorescent protein (GFP) (M0802-3a, Abiocode, Agoura Hills, CA, USA) (1:10,000), rabbit polyclonal anti-flag epitope (FLAG) (F7425, Sigma, St. Louis, MO, USA) (1:10,000), goat polyclonal anti-mouse coupled to horseradish peroxidase (A2554, Sigma, St. Louis, MO, USA) (1:15,000), and goat polyclonal anti-rabbit coupled to horseradish peroxidase (A0545, Sigma, St. Louis, MO, USA) (1:15,000).

### Bimolecular Fluorescence Complementation (BiFC)

Fully expanded young leaves of 4-week-old *N. benthamiana* plants were co-infiltrated with *A. tumefaciens* clones carrying the appropriate BiFC plasmids using a 1 mL needleless syringe and imaged two days post-infiltration with a Leica TCS SMD confocal microscope (Leica Microsystems) using the preset settings for YFP (Ex: 514 nm, Em: 525-575 nm). For nuclei staining, leaves were infiltrated with 5 μg/mL Hoechst 33258 (Sigma) solution and incubated in the dark for 30-60 minutes before observation by using the corresponding preset settings (Ex: 355 nm, Em: 430-480 nm).

### Yeast two-hybrid

pGBKT7-and pGADT7-based constructs were co-transformed into the Y2HGold yeast strain (Clontech) using Yeastmaker™ Yeast Transformation System 2 (Clontech) according to the manufacturer’s instructions. The co-transformants were selected on minimal synthetic defined (SD) media without leucine and tryptophan; interactions were tested on SD media without leucine, tryptophan, histidine, and adenine. pGADT7-T and pGBKT7-p53 constructs were used as positive control; empty vectors were used as negative control.

### Split-luciferase complementation imaging assay

*A. tumefaciens* strains carrying the appropriate plasmids were agroinfiltrated into 4-week-old *N. benthamiana* plants using a 1 mL needleless syringe. Two days post-infiltration, the same leaves were infiltrated with 1 mM D-luciferin solution and kept in the dark for 5 min before imaging. The luminescence images were captured using a CCD camera (NightShade LB 985, Berthold).

### Visualization of protein subcellular localization

For subcellular localization, plant tissues expressing GFP-or RFP-fused proteins were imaged with a Leica TCS SP8 confocal microscope (Leica Microsystems) using the preset settings for GFP (Ex: 488 nm, Em: 500-550 nm) or RFP (Ex: 554 nm, Em: 580-630 nm). Confocal imaging for co-localization of C2-GFP and TYLCV proteins fused to RFP was performed on a Leica TCS SP8 point scanning confocal microscope using the pre-set sequential scan settings for GFP (Ex:488 nm, Em:500–550 nm) and RFP (Ex:561 nm, Em:600–650 nm).

### TYLCV infection

For TYLCV local infection assays, fully expanded young leaves of 4-week-old *N. benthamiana* plants were infiltrated with *A. tumefaciens* carrying the TYLCV infectious clone (WT or mutants). Samples were collected at 2.5 days post-inoculation (dpi) to detect viral accumulation.

For TYLCV systemic infection assays, *A. tumefaciens* carrying the TYLCV infectious clone (WT or mutants) were syringe-inoculated in the stem of 2-week-old *N. benthamiana* plants. Leaf discs from the three youngest apical leaves were harvested at 21 dpi to detect viral accumulation.

### Determination of viral accumulation by quantitative PCR (qPCR)

To determine viral accumulation, total DNA was extracted from *N. benthamiana* leaves using the CTAB method (Minas et al., 2011).The DNA from local infection assays was treated with *Dpn*I at 37°C for 1 hour prior to further analysis. Quantitative PCR (qPCR) was performed with primers to amplify Rep (Wang et al., 2017b). The qPCR reaction was performed with Hieff® qPCR SYBR® Green Master Mix (Yeasen), with the following program: 3 min at 95°C, and 40 cycles consisting of 15 s at 95°C, 30 s at 60°C. As internal reference for DNA detection, the 25S ribosomal DNA interspacer (ITS) was used (Mason et al., 2008). qPCR was performed in a BioRad CFX96 real-time system as described previously (Wang et al., 2017b). The primers used are described in Supplemental table 8.

### Reverse transcription quantitative PCR (RT-qPCR)

RNA was extracted using the Plant RNA kit (OMEGA Bio-Tek); cDNA was prepared using the iScript™ gDNA Clear cDNA Synthesis Kit (Bio-Rad) according to the manufacturer’s instructions. The qPCR reaction was performed with Hieff® qPCR SYBR® Green Master Mix (Yeasen), with the following program: 3 min at 95°C, and 40 cycles consisting of 15 s at 95°C, 30 s at 60°C. *Elongation factor-1 alpha* (*NbEF1α*) (Nicot et al., 2005) or *Actin2 (NbACT)* (Viczián et al., 2014) were used as reference genes, as indicated. The primers used are described in Supplemental table 8.

### Bacterial infections

Four-week-old *N. benthamiana* leaves were infiltrated with a *P. syringae* pv. *tomato* DC3000 *ΔhopQ1-1* suspension (Rufian et al., 2018) (OD_600_ = 0.0002 in 10 mM MgCl_2_) using a 1 mL needleless syringe upon transient expression of the construct of interest. Bacterial growth was determined three days after inoculation by plating 1:10 serial dilutions of leaf extracts on solid LB medium (1% tryptone, 0.5% yeast extract, and 0.5% NaCl) with the appropriate antibiotics; plates were incubated at 28°C for two days before bacterial colony-forming units (cfu) were counted.

### RNA seq and analysis

Transcriptome sequencing in *N. benthamiana* was performed as previously described (Wu et al., 2019). Four biological replicates were used per sample. The paired-end reads were cleaned by Trimimomatic (Bolger et al., 2014) (version 0.36). Clean read pairs were retained for further analysis after trimming the adapter sequence, removing low quality bases, and filtering short reads. The *N. benthamiana* draft genome sequence (v1.0.1) (Bombarely et al., 2012) was downloaded from the Sol Genomics Network (ftp://ftp.solgenomics.net/genomes/*Nicotiana_benthamiana/assemblies/*). Clean reads were mapped to the genome sequence by HISAT (Kim et al., 2015) (version 2.1.0) with default parameters. The number of reads that were mapped to each *N. benthamiana* gene was calculated with the htseq-count script in HTSeq (Bombarely et al., 2012). Differentially expressed genes (DEGs) with at least 1.5 fold change in expression and a FDR < 0.05 between control and experiment samples were identified by using EdgeR (Robinson et al., 2010).

The heatmap with hierarchical clustering was drawn by R package pheatmap. Venn diagrams were drawn by Venny (http://bioinformatics.psb.ugent.be/webtools/Venn/) and modified in Adobe Illustrator. The *Arabidopsis thaliana* homologous genes of the DEGs identified in *N. benthamiana* were used for Gene Ontology (GO) term enrichment analysis in AgriGO v2.0 (Tian et al., 2017).

## Supporting information

Supplementary table 2. DEGs in RNA-seq experiments (from Figures 3 and 4).

Supplementary table 3. Functional enrichment analyses (from Figures 3D and 3E).

Supplementary table 4. Functional enrichment analysis (from Figure 4E).

Supplementary table 5. Functional enrichment analysis (from Supplementary figure 10)

Supplementary table 6. Primers used for cloning in this work

Supplementary table 7. Plasmids used in this work

Supplementary table 8. Primers used for qPCR and qRT-PCR in this work

## ACKNOWLEDGEMENTS

The authors thank past and present members of the Lozano-Duran lab for fruitful discussions; Xinyu Jian, Aurora Luque, the PSC Core Cell Biology Facility, and the PSC Core Genomics Facility for technical assistance; and Alberto P Macho for critical reading of this manuscript. This work was supported by the Strategic Priority Research Program of the Chinese Academy of Sciences (CAS) (grant number XDB27040206) and the Shanghai Center for Plant Stress Biology, CAS. RL-D is the recipient of a National Foreign Talents project (grant number G20200113006). LM-P is the recipient of a Young Investigator Grant from the Natural Science Foundation of China (NSFC) (grant number 31850410467), a President’s International Fellowship Initiative (PIFI) postdoctoral fellowship (2018PB058 and 2020PB0080) from CAS, and a Foreign Youth Talent Program project (grant number 20WZ2503900) from the Shanghai Science and Technology Commission. BGG is the recipient of a President’s International Fellowship Initiative (PIFI) postdoctoral fellowship (2020PB0082), a Talent-Introduction grant from the Chinese Postdoctoral International Exchange, and a Foreign Youth Talent Program project (grant number 20WZ2504500) from the Shanghai Science and Technology Commission. EA is the recipient of a Young Investigator Grant from the NSFC (grant number 31950410534), a Marie Skłodowska-Curie Grant from the European Union’s Horizon 2020 Research and Innovation Program (Grant 896910-GeminiDECODER), and a National Foreign Talents project (grant number QN20200113001).

