## Supplementary material for "Combinatorial interactions between viral proteins expand the functional landscape of the viral proteome"

[1]Shanghai Center for Plant Stress Biology, CAS Center for Excellence in Molecular Plant Sciences, Chinese Academy of Sciences, Shanghai 201602, China. [2]University of the Chinese Academy of Sciences, Beijing 100049, China. [3]Instituto de Hortofruticultura Subtropical y Mediterránea "La Mayora" (IHSM-UMA-CSIC), Area de Genética, Facultad de Ciencias, Universidad de Málaga, Campus de Teatinos s/n, E-29071 Málaga, Spain. [4]Department of Plant Biochemistry, Centre for Plant Molecular Biology (ZMBP), Eberhard Karls University, D-72076 Tübingen, Germany.

*These authors contributed equally to this work

[#]Corresponding author

### The supplementary material for this manuscript includes:

**Supplementary figure 1-10**

**Supplementary tables 1-8**

### SUPPLEMENTARY FIGURES

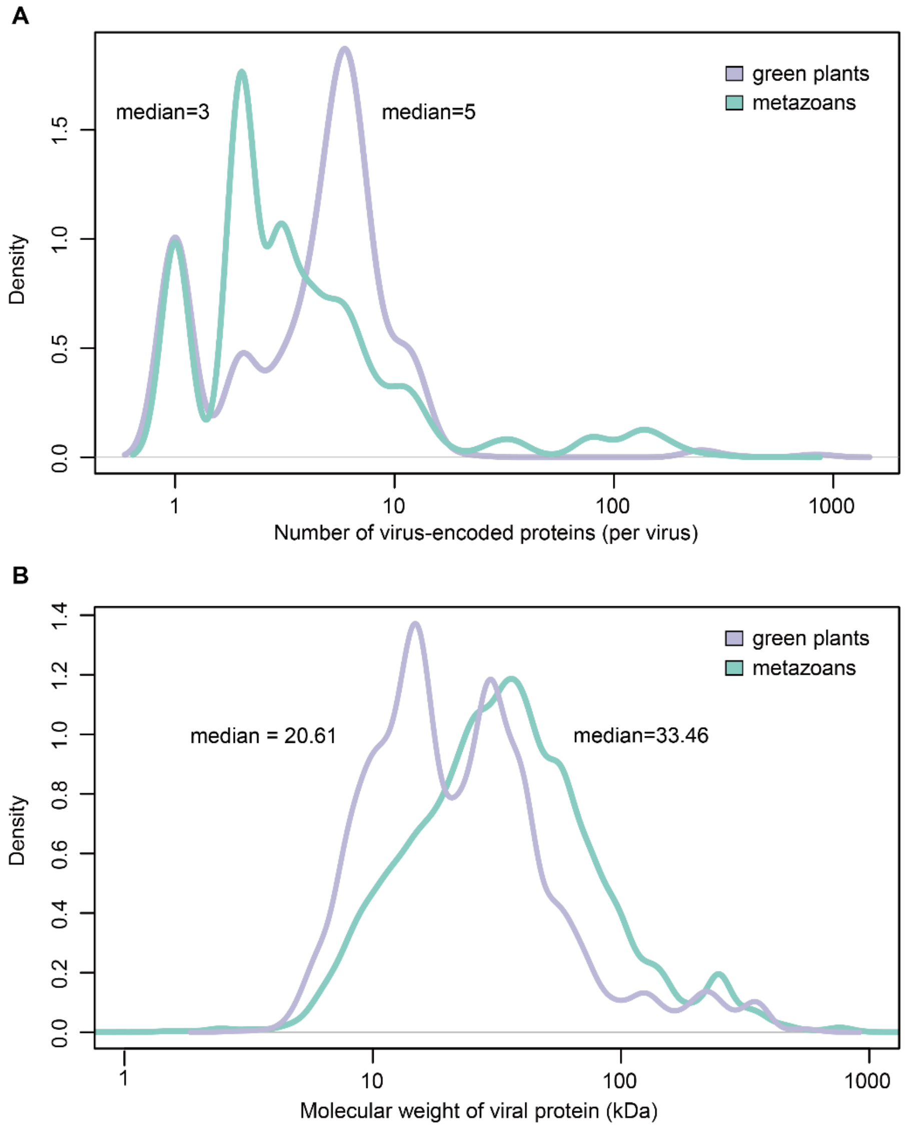


**Supplementary figure 1.** Average numbers of virus-encoded proteins and their molecular weight in animal and plant viruses. Sequences were downloaded from NCBI Virus, from complete RefSeq genome sequences of viruses infecting *Viridiplantae* (green plants, taxid: 33090) or *Metazoa* (metazoans, taxid: 33208).

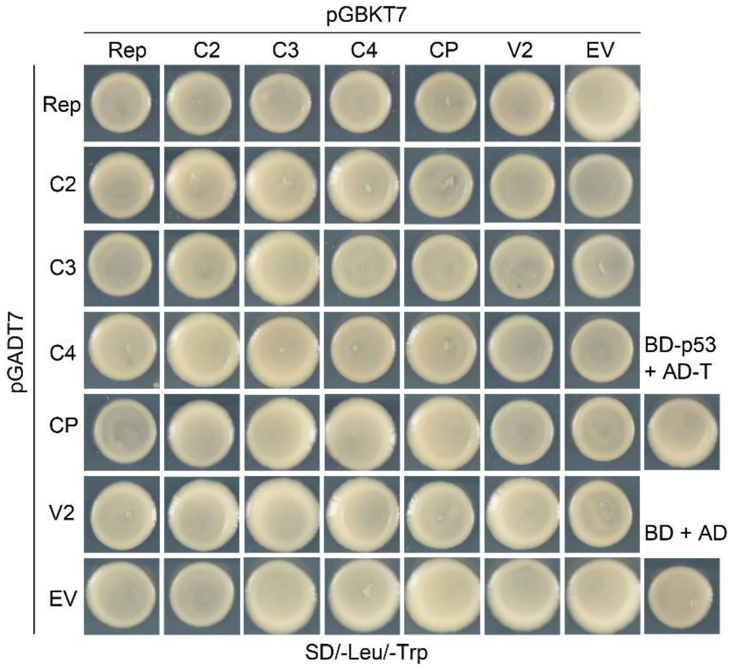


**Supplementary figure 2.** Yeast two-hybrid co-transformation control. The minimal synthetic defined (SD) medium without leucine (Leu) and tryptophan (Trp) was used to select co-transformants.

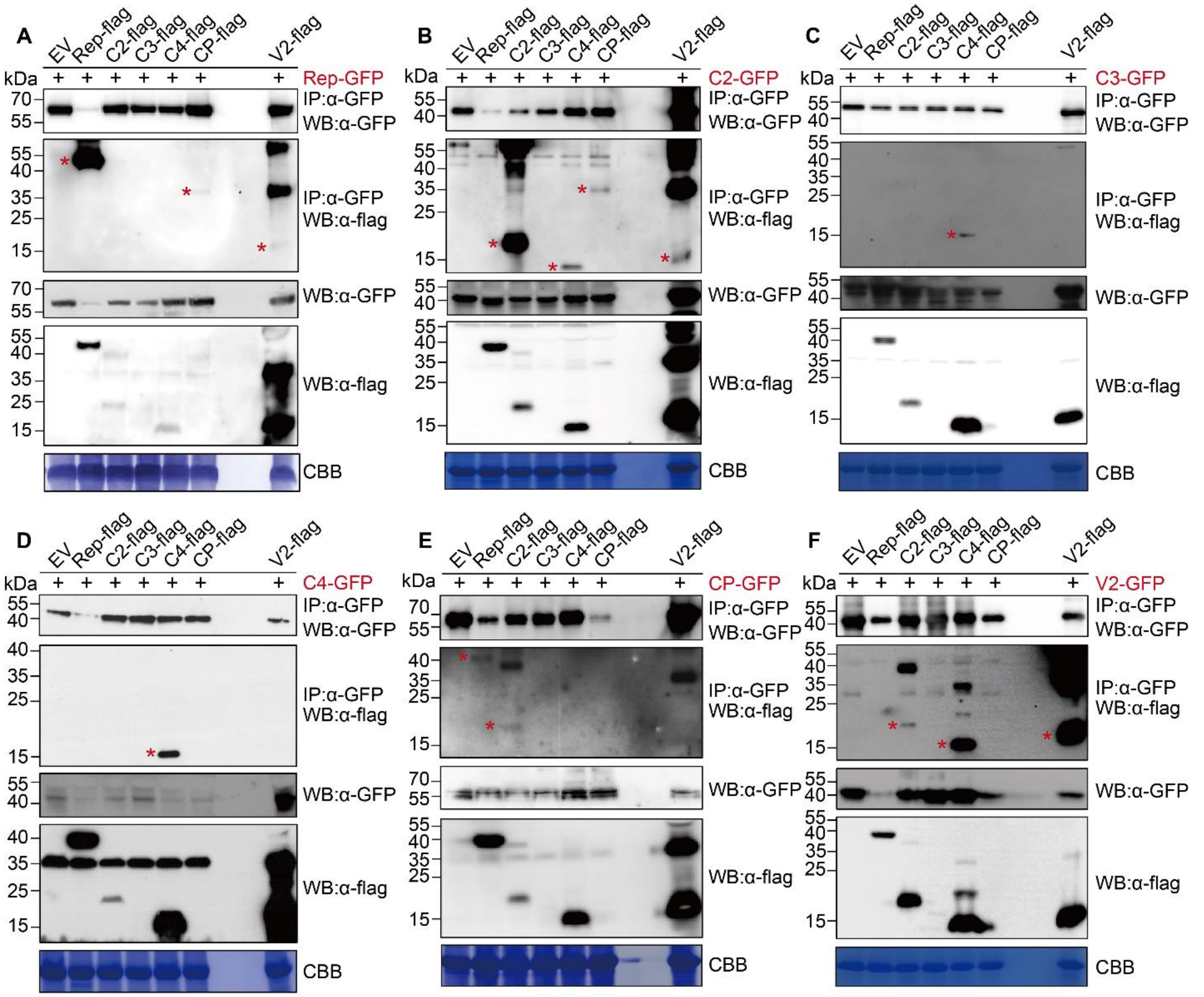


**G** Without TYLCV

| | Rep-GFP | C2-GFP | C3-GFP | C4-GFP | CP-GFP | V2-GFP |
|---|---|---|---|---|---|---|
| Rep-flag | +++ | --- | +-- | --- | +-- | +-- |
| C2-flag | --- | +++ | --- | --- | +-- | +-- |
| C3-flag | --- | --- | --- | --- | --- | --- |
| C4-flag | --- | ++- | ++- | +++ | --- | +++ |
| CP-flag | +-- | ++- | --- | --- | --- | --- |
| V2-flag | +-- | ++- | --- | --- | --- | +++ |

**Supplementary figure 3.** Representative co-immunoprecipitation (co-IP) assays of Rep-, C2-, C3-, C4-, CP- and V2-flag with Rep- (**A**), C2- (**B**), C3- (**C**), C4- (**D**), CP- (**E**) or V2-GFP (**F**) following transient expression in *N. benthamiana* leaves. IB: immunoblotting, IP: immunoprecipitation, CBB: Coomassie brilliant blue. Molecular weight of Rep-, C2-, C3-, C4-, CP-, and V2-GFP is 65, 42, 43, 38, 57 and 40 kDa, respectively; molecular weight of Rep-, C2-, C3-, C4-, CP-, and V2-flag are 41, 15, 16, 11, 30 and 14 kDa, respectively. Asterisks indicate the expected band for each protein. (**G)** Summary table containing the results of all co-IP replicates performed in the absence of the virus. Column headings indicate the viral protein used as a bait; row headings indicate prey proteins.

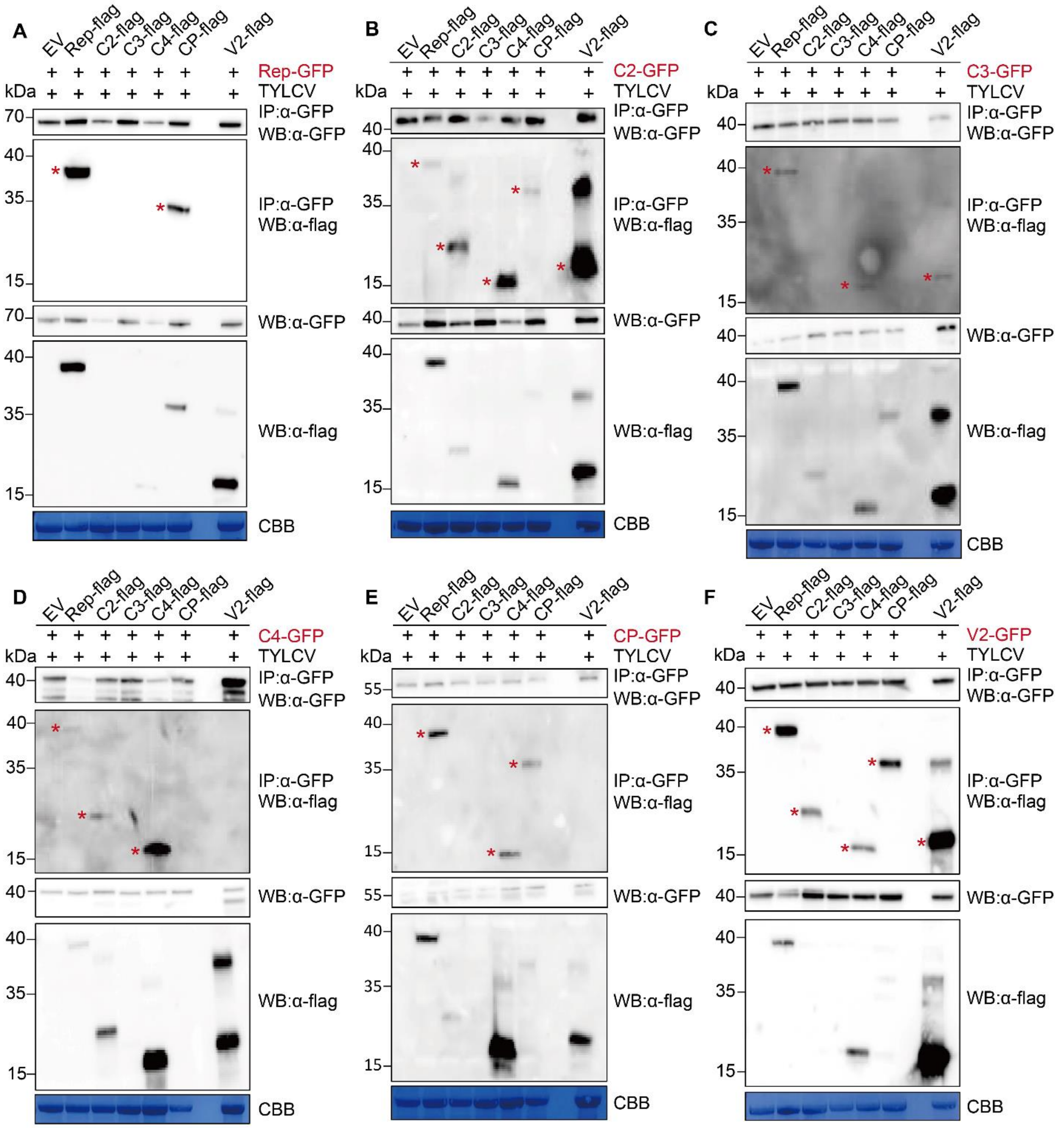


**G** With TYLCV

| | REP-GFP | C2-GFP | C3-GFP | C4-GFP | CP-GFP | V2-GFP |
|---|---|---|---|---|---|---|
| REP-flag | ++ | --++ | -+++ | -+-+ | -+++ | ++ |
| C2-flag | -- | ++++ | ---- | --++ | ---- | ++ |
| C3-flag | -- | ---- | -+-- | ---- | ---- | -- |
| C4-flag | -- | --++ | -+++ | -+++ | -+-+ | ++ |
| CP-flag | ++ | --++ | -+-- | -+-- | +-++ | ++ |
| V2-flag | -- | -+++ | -+++ | -+-- | --+- | ++ |

**Supplementary figure 4.** Representative co-immunoprecipitation (co-IP) assays of Rep-, C2-, C3-, C4-, CP- and V2-flag with Rep- (**A**), C2- (**B**), C3- (**C**), C4- (**D**), CP- (**E**) or V2-GFP (**F**) following transient expression in *N. benthamiana* leaves in the presence of the virus. IB: immunoblotting, IP: immunoprecipitation, CBB: Coomassie brilliant blue. Molecular weight of Rep-, C2-, C3-, C4-, CP- and V2-GFP is 65, 42, 43, 38, 57 and 40 kDa, respectively; molecular weight of Rep-, C2-, C3-, C4-, CP- and V2-flag are 41, 15, 16, 11, 30 and 14 kDa, respectively. Asterisks indicate the expected band for each protein. (**G)** Summary table containing the results of all co-IP replicates performed in the presence of the virus. Column headings indicate the viral protein used as a bait; row headings indicate prey proteins.

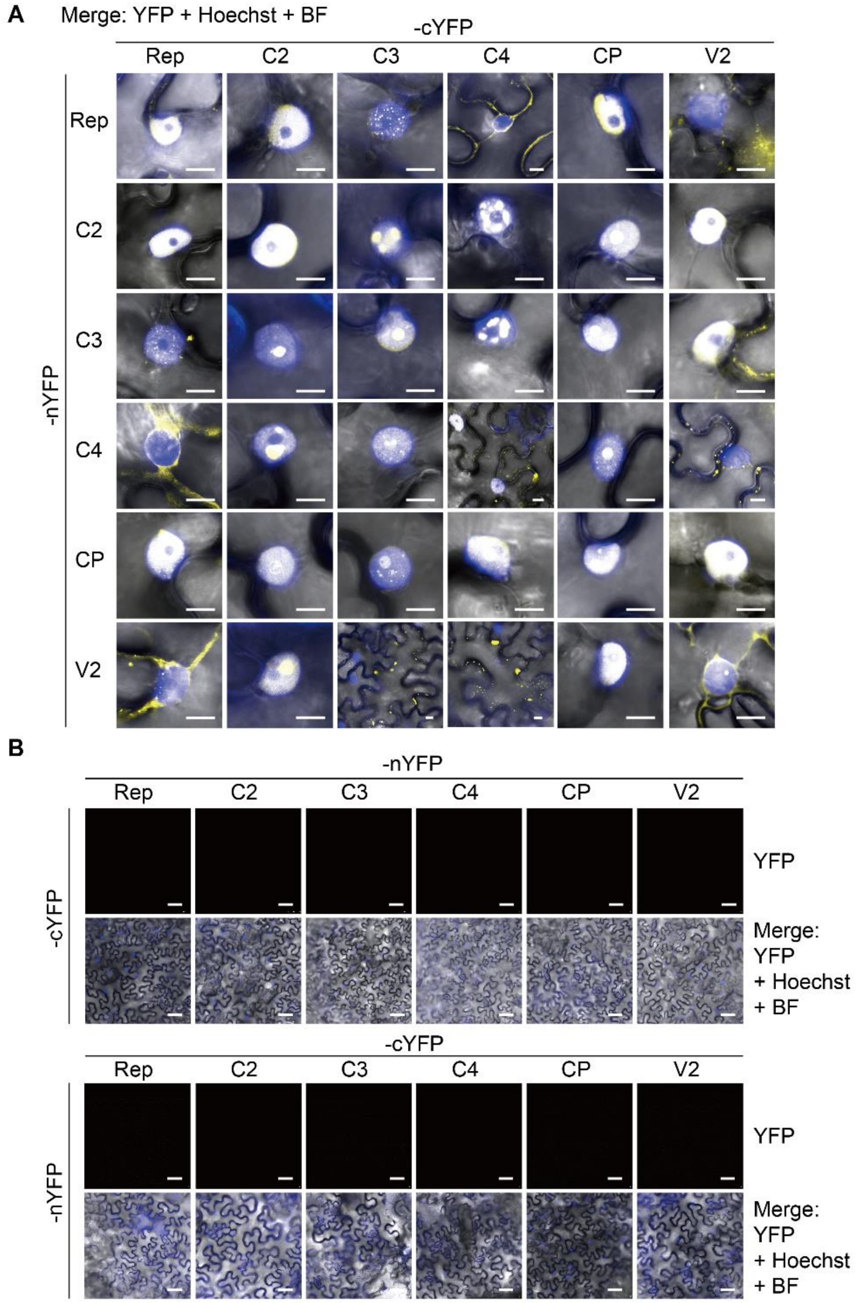


**Supplementary figure 5.** Viral protein-protein interactions detected by bimolecular fluorescence complementation (BiFC) in *N. benthamiana* leaves combined with Hoechst staining (**A**) and negative controls (**B**). nYFP: N-terminal half of the YFP; cYFP: C-terminal half of the YFP; BF: bright field. Images were taken at 2 days post-infiltration (dpi). Scale bar = 10 µm in (A) or 50 µm in (B). This experiment was repeated at least four times with similar results.

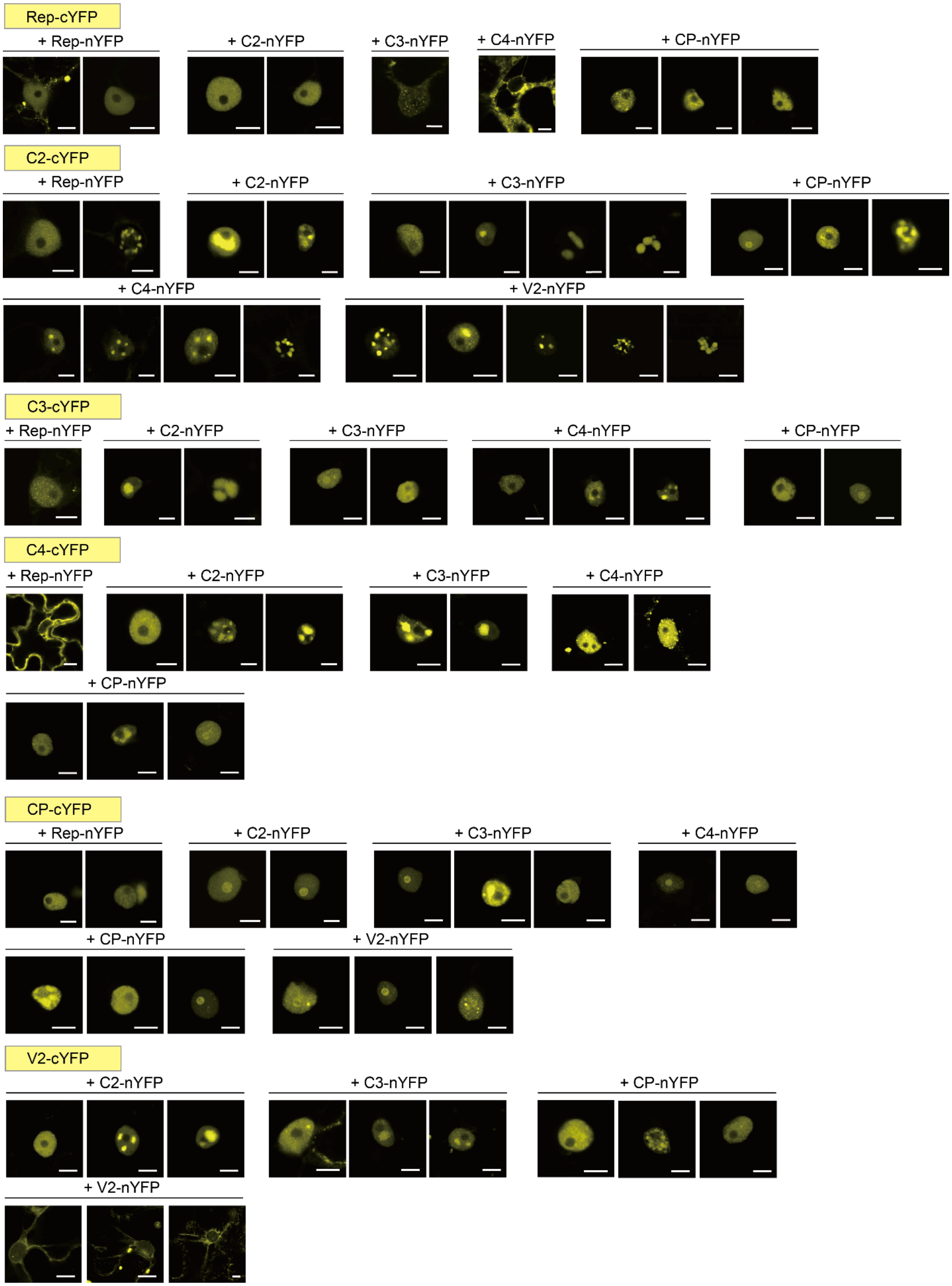


**Supplementary figure 6.** Additional BiFC images (from Figure 1C). nYFP: N-terminal half of the YFP; cYFP: C-terminal half of the YFP. Images were taken at 2 days post-infiltration (dpi). Scale bar = 10 µm. This experiment was repeated at least four times with similar results.

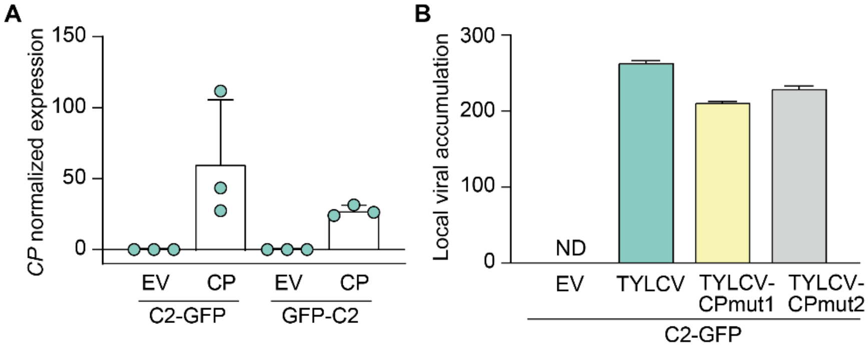


**Supplementary figure 7.** Accumulation of the CP transcript and viral accumulation in the samples from Figure 2C and 2D, respectively. (**A**) Accumulation of the CP transcript (from Figure 2C), measured by qRT-PCR. *NbEF1α* was used as the normalizer. Values represent the mean of three plants. Error bars represent SD. (**B**) Accumulation of viral DNA in samples from Figure 2D. *ITS* was used as the normalizer. Values represent the mean of three plants. Error bars represent SD.

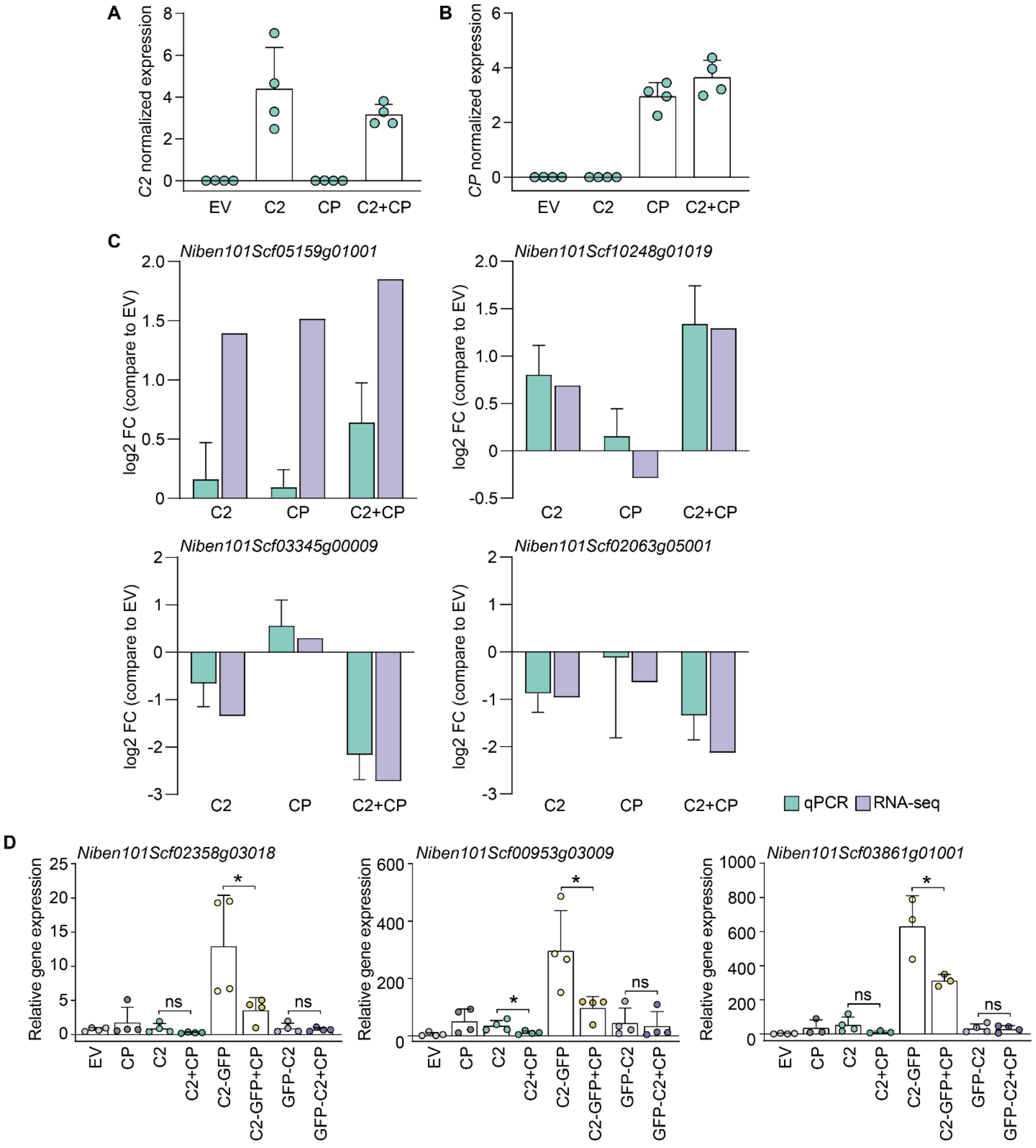


**Supplementary figure 8.** Validation of the expression of plant and viral genes in the samples used for RNA-seq (Figure 3). (**A** and **B**) C2 (A) and CP (B) transcript accumulation, measured by qRT-PCR. Expression values are relative to *NbACT*. Results are the mean of four biological replicates. Error bars represent SD. (**C**) Comparison of the accumulation of transcripts of selected DEGs in the RNA-seq data and as measured by qRT-PCR. Expression values are the mean of log2 FC, relative to EV, from four biological replicates. *NbACT* was used as the normalizer. FC: fold change; EV: empty vector. (**D**) Expression of selected DEGs upon expression of C2, C2-GFP, GFP-C2 in the presence and absence of CP in *N. benthamiana* leaves. Samples expressing CP or empty vector (EV) are used as control. Expression values are the mean of at least three biological replicates. Error bars represent SD. Asterisks indicate a statistically significant difference (*: $p<0.05$) according to a two-tailed comparison t-test. *NbACT* was used as the normalizer.

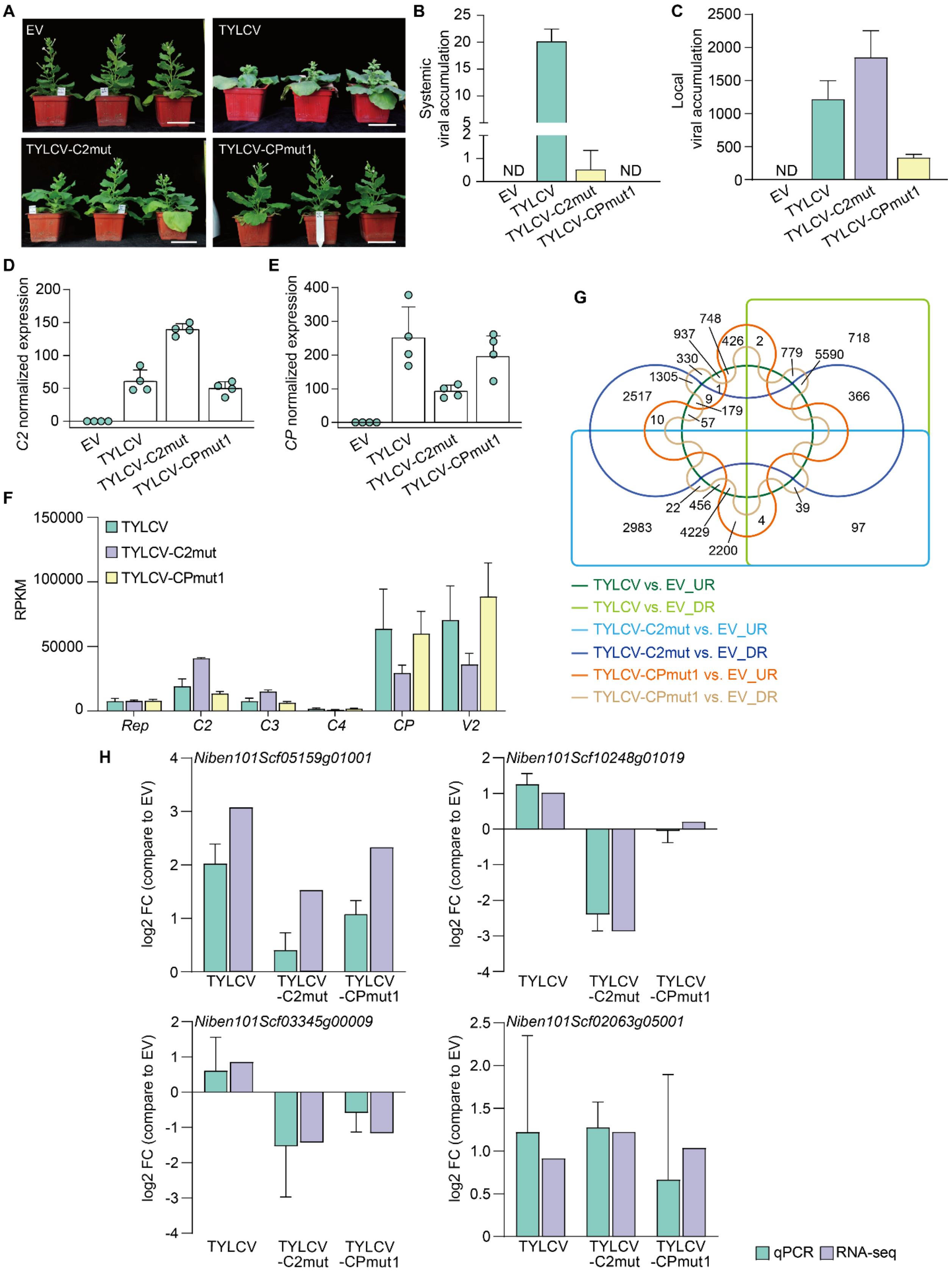

A
EV
TYLCV
TYLCV-C2mut
TYLCV-CPmut1
B
Systemic viral accumulation
ND
ND
EV
TYLCV
TYLCV-C2mut
TYLCV-CPmut1
C
Local viral accumulation
ND
D
C2 normalized expression
E
CP normalized expression
F
RPKM
TYLCV
TYLCV-C2mut
TYLCV-CPmut1
Rep
C2
C3
C4
CP
V2
G
TYLCV vs. EV_UR
TYLCV vs. EV_DR
TYLCV-C2mut vs. EV_UR
TYLCV-C2mut vs. EV_DR
TYLCV-CPmut1 vs. EV_UR
TYLCV-CPmut1 vs. EV_DR
H
Niben101Scf05159g01001
Niben101Scf10248g01019
Niben101Scf03345g00009
Niben101Scf02063g05001
log2 FC (compare to EV)
TYLCV
TYLCV -C2mut
TYLCV -CPmut1
qPCR
RNA-seq

**Supplementary figure 9.** Validation of the expression of plant and viral genes in the samples used for RNA-seq (Figure 4). (**A)** Symptoms in *N. benthamiana* plants inoculated with TYLCV WT or C2-/CP-null mutants (TYLCV-C2mut and TYLCV-CPmut1, respectively), or inoculated with empty vector (EV) as negative control. Pictures were taken at 21 days post-inoculation (dpi). Scale bar: 10 cm. (**B)** Viral DNA accumulation in systemic infections in *N. benthamiana* plants measured by qPCR. Values are the mean of three independent biological replicates. Error bars represent SD. Samples were taken at 21 dpi. *ITS* was used as the normalizer. (**C**) Viral DNA accumulation in *N. benthamiana* leaves infiltrated with TYLCV WT or C2-/CP-null mutants (TYLCV-C2mut and TYLCV-CPmut1, respectively), or transformed with empty vector (EV) as negative control. Samples were taken at 2.5 days post-inoculation (dpi). *ITS* was used as the normalizer. Values represent the mean of six plants. Error bars represent SD. (**D** and **E**) C2 (D) and CP (E) transcript accumulation. Expression values are relative to *NbACT*. Results are the mean of four biological replicates. Error bars represent SD. (**F**) Expression of TYLCV genes in the different samples as detected by RNA-seq. RPKM: reads per kilobase of transcript per million mapped reads. (**G**) Venn diagram of the subsets of up- and down-regulated genes in the samples infected with TYLCV WT or C2-/CP-null mutants (TYLCV-C2mut and TYLCV-CPmut1) compared to the empty vector control (EV). UR: up-regulated; DR: down-regulated. (**H**) Expression of selected DEGs. Expression values are the mean of log2 FC, relative to samples inoculated with the EV, from four biological replicates. *NbACT* was used as the normalizer. FC: fold change; EV: empty vector.

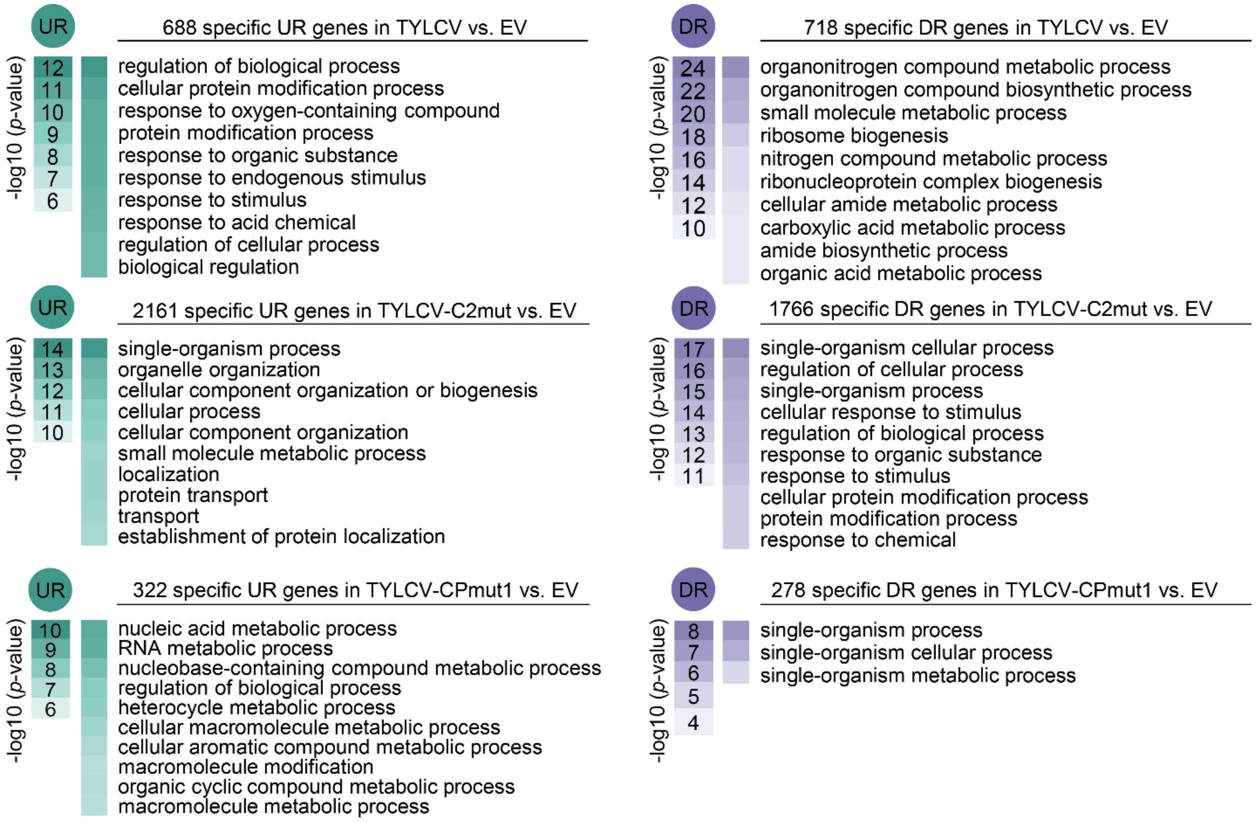


**Supplementary figure 10.** Functional enrichment analysis of the subsets of up-regulated (UR) or down-regulated (DR) genes in Figure 4A. Gene Ontology (GO) categories from the Biological Process ontology enriched with a *p*-value<0.01 (up to top 10) are shown; functional enrichment analysis was performed using the orthologues in *A. thaliana*. The colour scale indicates the -log10 (*p*-value), showing the significance of GO term enrichment. For a full list, see Supplementary table 5.

- **SUPPLEMENTARY TABLES**

**Supplementary table 1.** AP-MS/MS data.

**Supplementary table 2.** DEGs in RNA-seq experiments (from Figures 3 and 4).

**Supplementary table 3.** Functional enrichment analysis (from Figures 3D and 3E).

**Supplementary table 4.** Functional enrichment analysis (from Figure 4E).

**Supplementary table 5.** Functional enrichment analysis (from Supplementary figure 10).

**Supplementary table 6.** Primers used for cloning in this work.

**Supplementary table 7.** Plasmids used in this work.

**Supplementary table 8.** Primers used for qPCR and qRT-PCR in this work.
